# Spindle asymmetry drives non-Mendelian chromosome segregation

**DOI:** 10.1101/180869

**Authors:** Takashi Akera, Lukáš Chmátal, Emily Trimm, Karren Yang, Chanat Aonbangkhen, David M. Chenoweth, Carsten Janke, Richard M. Schultz, Michael A. Lampson

**Affiliations:** Department of Biology, School of Arts and Sciences, University of Pennsylvania, Philadelphia, Pennsylvania 19104, USA; Department of Chemistry, School of Arts and Sciences, University of Pennsylvania, Philadelphia, Pennsylvania 19104, USA; Institut Curie, PSL Research University, CNRS UMR3348, Centre Universitaire, Bâtiment 110, F-91405 Orsay, France

## Abstract

Genetic elements compete for transmission through meiosis, when haploid gametes are created from a diploid parent. Selfish elements can enhance their transmission through meiotic drive, in violation of Mendel’s Law of Segregation. In female meiosis, selfish elements drive by preferentially attaching to the egg side of the spindle, which implies some asymmetry between the two sides of the spindle, but molecular mechanisms underlying spindle asymmetry are unknown. Here we show that CDC42 signaling from the cell cortex regulates microtubule tyrosination to induce spindle asymmetry, and non-Mendelian segregation depends on this asymmetry. These signals depend on cortical polarization directed by chromosomes, which are positioned near the cortex to allow the asymmetric cell division. Thus, selfish meiotic drivers exploit the asymmetry inherent in female meiosis to bias their transmission.

---

Genetic conflict is inherent in any haploid-diploid life cycle because genetic elements compete for transmission to the offspring through meiosis, the process by which haploids are generated. Mendel’s Law of Segregation states that alleles of a gene are transmitted with equal probability, but it is increasingly clear that this law is often violated, and segregation can be manipulated by selfish genetic elements through meiotic drive. Drive can occur by eliminating competing gametes that do not contain the selfish element (e.g., sperm killing or spore killing) or by exploiting the asymmetry in female meiosis to increase the transmission of the selfish element to the egg. Although the impact of meiotic drive on many aspects of evolution and genetics is now recognized, with examples widespread across eukaryotes (*1*–*4*), the underlying mechanisms are largely unknown.

Female meiosis provides a clear opportunity for selfish elements to cheat because of its inherent asymmetry: only chromosomes that segregate to the egg can be transmitted to offspring, while the rest are degraded in polar bodies. Conceptually, female meiotic drive depends on three conditions: asymmetry in cell fate, a functional difference between homologous chromosomes that influences their segregation, and asymmetry within the meiotic spindle (*5*). The asymmetry in cell fate is well established (*6*), and chromosomal rearrangements and amplifications of repetitive sequences (e.g., centromeres) are associated with biased segregation (*7*–*10*). Asymmetry within the meiotic spindle was noted in grasshopper in 1976 (*11*), but not studied further, and molecular mechanisms regulating such asymmetry are unknown.

Oocyte spindles are positioned close to the cortex and oriented perpendicular to the cortex in order to achieve the highly asymmetric cell division, so that cytokinesis produces a large egg and a small polar body (Fig. 1A). A selfish element drives by preferentially attaching to the egg side of the spindle, implying some difference in microtubules (MTs) between the egg side and the cortical side. To determine how such spindle asymmetry is regulated, we first screened for a marker for asymmetry using mouse oocytes as a model in which we have observed meiotic drive (*10*, *12*). MTs can be functionally diversified by post-translational modifications (PTMs, Fig. S1A) (*13*–*15*), and we tested for asymmetry in these modifications. Among the PTMs that we examined, tyrosinated (Tyr) and detyrosinated (dTyr) α-tubulin showed complementary asymmetry on the spindle, with the cortical side enriched for Tyr α-tubulin and the egg side for dTyr α-tubulin (Fig. 1, B and D). ß-tubulin, acetylated α-tubulin, and poly-glutamylated tubulin did not show significant asymmetry (Fig. 1D; Fig. S1B).

**Fig. 1.**
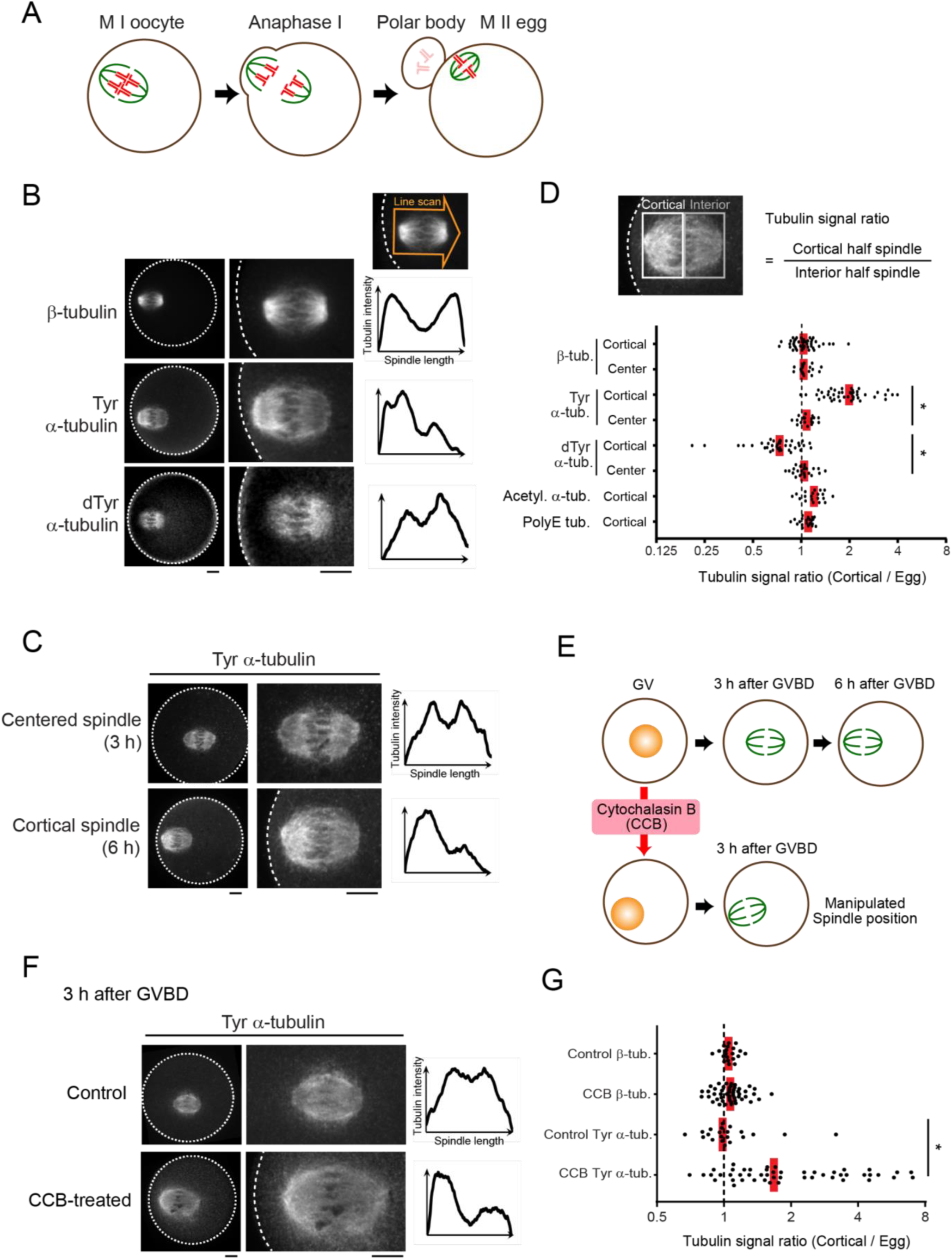
Cortical proximity induces asymmetry within the mouse oocyte spindle. (**A**) Schematic of asymmetric female meiosis I in mouse and spindle orientation perpendicular to the cortex. (**B-G**) CF-1 oocytes were fixed at metaphase I and stained for the indicated post-translational modifications on tubulin. Cortical spindles (B-D) were examined at 6 h after GVBD, and centered spindles (C, D) at 3 h GVBD. Treatment with cytochalasin B generates cortical spindles at 3 h after GVBD in 24 % of oocytes, and asymmetry was measured in these oocytes at this time (E-G). Images (B, C, F) are sum intensity z-projections showing the whole oocyte (left) or a magnified view of the spindle (right), with the dashed line indicating the cortex, and graphs are line scans of tubulin intensity across the spindle. Spindle asymmetry was quantified (D, G) as the ratio of the cortical half to the interior half (n > 18 spindles for each sample). Each dot represents a single spindle; red line, median; *p < 0.0001. Scale bars, 10 μm.

As a clue to how spindle asymmetry might be established, we found that spindles were asymmetric late in metaphase I when positioned near the cortex, but not earlier when positioned in the center of the oocyte (Fig. 1, C to E; Fig. S2). Because the MI spindle first forms in the center and then migrates towards the cortex (*16*–*20*), asymmetry might depend on either cortical proximity or time, or both. To distinguish between these possibilities, we manipulated spindle position by treating oocytes with cytochalasin B (CCB) before maturation. This treatment inhibits actin polymerization and leads to the nucleus drifting to the cortex in 24% of oocytes (Fig. 1E; Fig. S3A). As a result, the spindle is positioned near the cortex by 3 h after germinal vesicle breakdown (GVBD), much earlier than the migration at 6 h under normal conditions (Fig. 1E). Cortical spindles in CCB-treated oocytes showed asymmetric Tyr α-tubulin staining at 3 h after GVBD, whereas ß-tubulin staining remained symmetric (Fig. 1, F and G; Fig. S3B). Similar results were obtained with cytochalasin D (Fig. S3C). These results indicate that cortical proximity directly induces spindle asymmetry. Because one spindle pole generally faces the cortex, one possible mechanism to create asymmetry is a difference between the spindle poles, with the cortical pole generating higher levels of Tyr α-tubulin. This mechanism is unlikely, however, since mis-oriented spindles that are parallel to the cortex have stronger Tyr α-tubulin signals on the cortical side, which cannot be explained by a difference between spindle poles (Fig. S4). Together, these results suggest that the cortex directly regulates MTs to induce asymmetry within the spindle.

The cortex overlying the spindle is polarized by the chromosomes, through a chromatin-based gradient of RAN^GTP^ (*21*, *22*)(Fig. 2A). This polarized cortex is enriched in multiple signaling factors, including active CDC42 and RAC GTPases, and in polymerized actin (called the actin cap) (*6*, *23*, *24*) (Fig. 2A). To determine whether spindle asymmetry depends on this pathway, we prevented polarization by expressing either constitutively-active (RAN^Q69L^) or dominant-negative (RAN^T24N^) RAN mutants. In each case, loss of cortical polarization led to loss of spindle asymmetry (Fig. 2, B and C; Fig. S5A).

**Fig. 2.**
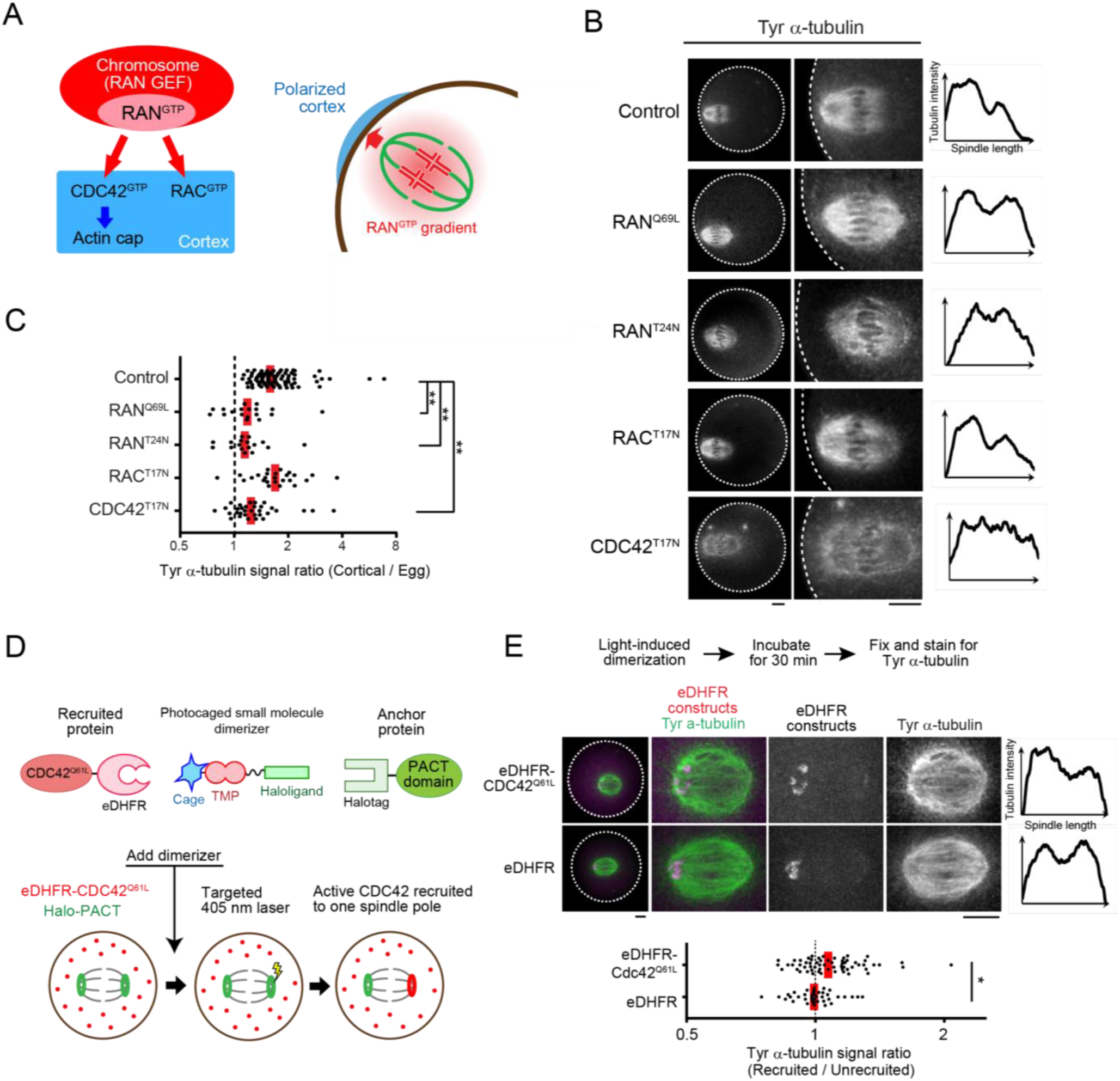
Cortical polarization and localized CDC42 signaling induce spindle asymmetry. (**A**) Schematic of cortical polarization. (**B, C**) CF-1 oocytes expressing the indicated GTPase mutant were fixed 6 h after GVBD and stained for Tyr α-tubulin. Images are sum intensity z-projections showing the whole oocyte (left) or a magnified view of the spindle (right), and graphs are line scans of tubulin intensity across the spindle. Spindle asymmetry was quantified (C) as the ratio of the cortical half to the interior half (n > 17 spindles for each sample). (**D**) Schematics of the light-induced dimerization system and experimental design. The small molecule dimerizer is composed of a Halo ligand linked to the eDHFR ligand Trimethoprim (TMP), which is photocaged. The PACT domain, tagged with EGFP and Halo, localizes to spindle poles, and CDC42^Q61L^ is tagged with mCherry and eDHFR. The dimerizer covalently binds Halo-PACT at spindle poles, and eDHFR-Cdc42^Q61L^ is recruited to one pole by local uncaging with light. (**E**) Halo-EGFP-PACT was co-expressed with either mCherry-eDHFR-Cdc42^Q61L^ (top) or mCherry-eDHFR (bottom) in CF-1 oocytes. Recruitment of eDHFR fusion proteins was induced by uncaging at one spindle pole. 30 min after uncaging, oocytes were fixed and stained for Tyr α-tubulin. Images are maximum intensity z-projection showing whole oocytes (left) or magnified views of the spindle, and graphs are line scans of tubulin intensity across the spindle. Spindle asymmetry was quantified as the ratio of the recruited side to the unrecruited side (n > 39 spindles for each sample). Each dot represents a single spindle; red line, median; *p < 0.01; **p < 0.0001. Scale bars, 10 μm.

To understand how the polarized cortex induces spindle asymmetry, we tested CDC42 and RAC GTPases. Expressing a dominant-negative CDC42 mutant (CDC42^T17N^) diminished the Tyr α-tubulin signal overall and prevented the asymmetry, whereas dominant-negative RAC mutant (RAC^T17N^) did not affect asymmetry (Fig. 2, B and C; Fig. S5, A and B). Furthermore, expressing a constitutively-active CDC42^Q61L^ mutant with the plasma membrane targeting CAAX motif removed (eDHFR-CDC42^Q61L^ΔCAAX) (*25*) significantly increased Tyr α-tubulin signal (Fig. S7). Because CDC42 activity is required for actin cap formation at the polarized cortex (*24*) (Fig. 2A), we tested whether the actin cap contributes to spindle asymmetry. Inhibiting the actin nucleating ARP2/3 complex, using the small molecule inhibitor CK-666, abolished actin cap formation as in CDC42^T17N^-expressing oocytes (*26*), but did not affect spindle asymmetry (Fig. S6). Together these findings demonstrate that active CDC42 GTPase is sufficient to increase α-tubulin tyrosination and required for spindle asymmetry independent of actin cap formation.

Our observations suggest that asymmetric localization of active CDC42 relative to the spindle is the mechanism underlying spindle asymmetry. To test this hypothesis, we developed an optogenetic strategy to target active CDC42 to one pole of a centered spindle, which is normally symmetric, using a photocaged small molecule that heterodimerizes Halotag and *E. coli* DHFR (eDHFR) fusion proteins (*27*, *28*) (Fig. 2D). We fused Halotag to the PACT domain of AKAP9, which localizes to spindle poles (*29*), and eDHFR to the constitutively-active CDC42^Q61L^ΔCAAX mutant. The dimerizer covalently binds to Halotag at spindle poles, and local uncaging with 405 nm light recruits eDHFR fusion proteins specifically to one spindle pole through Halotag-eDHFR dimerization (Fig. S8A). This recruitment happens within 2 min and lasts more than 30 min (Fig. S8A). Recruiting CDC42^Q61L^ΔCAAX to one spindle pole induced spindle asymmetry by increasing Tyr α-tubulin signals on the recruited side (Fig. 2E; Fig. S8B), whereas recruiting eDHFR alone had no effect. These results strongly support our model that cortically localized CDC42 activity induces asymmetry within the spindle. Several factors may contribute to the weaker asymmetry induced by our optogenetic approach, compared to the asymmetry observed normally on spindles near the cortex. First, Tyr α-tubulin is high overall in cells expressing CDC42^Q61L^ΔCAAX (Fig. S7), which leaves less opportunity to create asymmetry when both sides are high before dimerization. Second, experimentally induced levels of CDC42 at spindle poles may be lower than normal levels at the cortex, and finally other cortical factors may also contribute to the asymmetry.

To determine the significance of the observed spindle asymmetry for meiotic drive, we measured the biased orientation of selfish centromeres towards the egg pole (Fig. 3A). Previously, we showed that in a cross between two strains, CHPO and CF-1, bivalents in the hybrid oocytes have both “weaker” and “stronger” centromeres, inherited from CHPO and CF-1, respectively (*10*, *12*). Stronger centromeres have higher levels of both inner and outer kinetochore proteins and more minor satellite DNA containing binding sites for the centromere protein CENP-B. By expressing fluorescently-tagged CENP-B in these hybrid oocytes, we can distinguish stronger and weaker centromere in live cells. In this system, we showed that stronger centromeres preferentially orient towards the egg pole just before anaphase I (*10*) (Fig. 3B, late meta I). To directly test whether biased orientation depends on spindle asymmetry, which was also confirmed in this hybrid strain (Fig. S9), we abolished the asymmetry by expression of the RAN^Q69L^ or CDC42^T17N^ mutants. We find that the bias is lost under these conditions, demonstrating that meiotic drive depends on spindle asymmetry induced by cortical polarization (Fig. 3B).

**Fig. 3.**
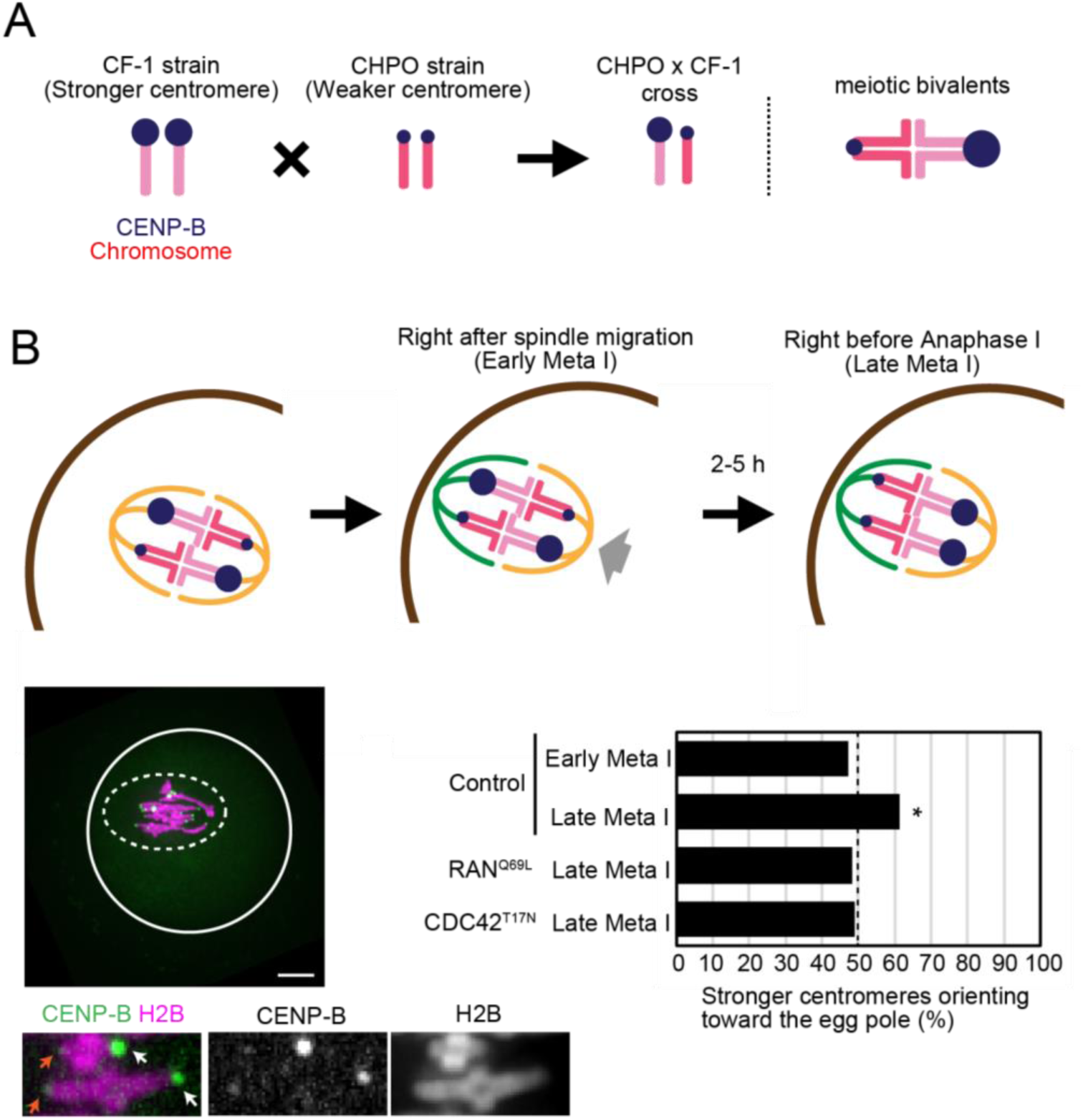
Spindle asymmetry is essential for biased orientation of selfish centromeres. (**A**) Schematic of biased orientation assay. A strain with stronger centromeres (CF-1) is crossed to a strain with weaker centromeres (CHPO). Bivalents in the hybrid offspring contain both stronger and weaker centromeres, which can be distinguished by CENP-B levels. (**B**) CHPO x CF-1 hybrid oocytes expressing CENP-B-EGFP and H2B-mCherry were imaged live, either shortly after spindle migration to the cortex (within 30 min, early meta I), or shortly before anaphase onset (within 30 min, late meta I). Image is a maximum intensity z-projection showing late meta I; white line: oocyte cortex, dashed line: spindle outline. Insets are optical slices showing two bivalents; arrows indicate stronger (white) and weaker (orange) centromeres; scale bar, 10 μm. The fraction of bivalents with the stronger centromere oriented towards the egg was quantified; n=152 bivalents for early meta I, 204 for late meta I (control), 108 for Ran^Q69L^ and 143 for CDC42^T17N^. * indicates significant deviation from 50% (p < 0.005).

Initial MT attachments are established before spindle migration to the cortex (*30*), while the spindle is symmetric, raising the question of whether biased orientation exists when spindles first migrate. We examined CHPO x CF-1 hybrid oocytes early in metaphase I, shortly after spindle migration. We do not find biased orientation at this stage (Fig. 3B, early meta I), indicating that the bias arises from re-orientation or flipping of stronger centromeres from the cortical to the egg side of the spindle while it is cortically positioned and asymmetric. CHPO x CF-1 hybrid oocytes remain in MI for 2-5 h after spindle migration to the cortex, likely due to chromosomes positioned off-center on the spindle (*12*, *31*) (Fig. 3B), which would provide time for these re-orientation events.

Consistent with this idea, we find several examples of bivalents flipping after spindle migration in hybrid oocytes (21 events in 23 cells) (Fig. 4A), as observed previously in a different strain (*30*). For these flipping events to establish biased orientation, they must preferentially occur in one direction, suggesting that one orientation is relatively more unstable than the other. This differential stability implies both a difference between centromeres of homologous chromosomes in how they interact with spindle MTs and a difference between the cortical and egg sides of the spindle. To test for these differences in hybrid oocytes, we examined kinetochore-MT fibers that remain stable at low temperature, while other MTs depolymerize (*32*). We find that stronger centromeres have more unstable attachments compared to weaker centromeres, and more dramatically when facing the cortical side of the spindle (Fig. 4B). These results show both that stronger centromeres are more likely to detach and that the cortical side is more susceptible to detachment. To test whether the enrichment of Tyr α-tubulin makes the cortical side more unstable, we manipulated tyrosination levels by modulating the expression of Tubulin Tyrosine Ligase (TTL), which catalyze α-tubulin tyrosination (*33*). Increasing Tyr α-tubulin by overexpressing TTL destabilized spindle MTs (Fig. 4C; Fig. S10A) based on sensitivity to low temperature (*34*). In contrast, decreasing Tyr α-tubulin by knocking down TTL stabilized spindle MTs (Fig. 4D; Fig. S10B). Collectively, these results indicate that the asymmetry in Tyr α-tubulin allows stronger centromeres to differentially interact with two sides of the spindle to preferentially orient towards the egg pole.

**Fig. 4.**
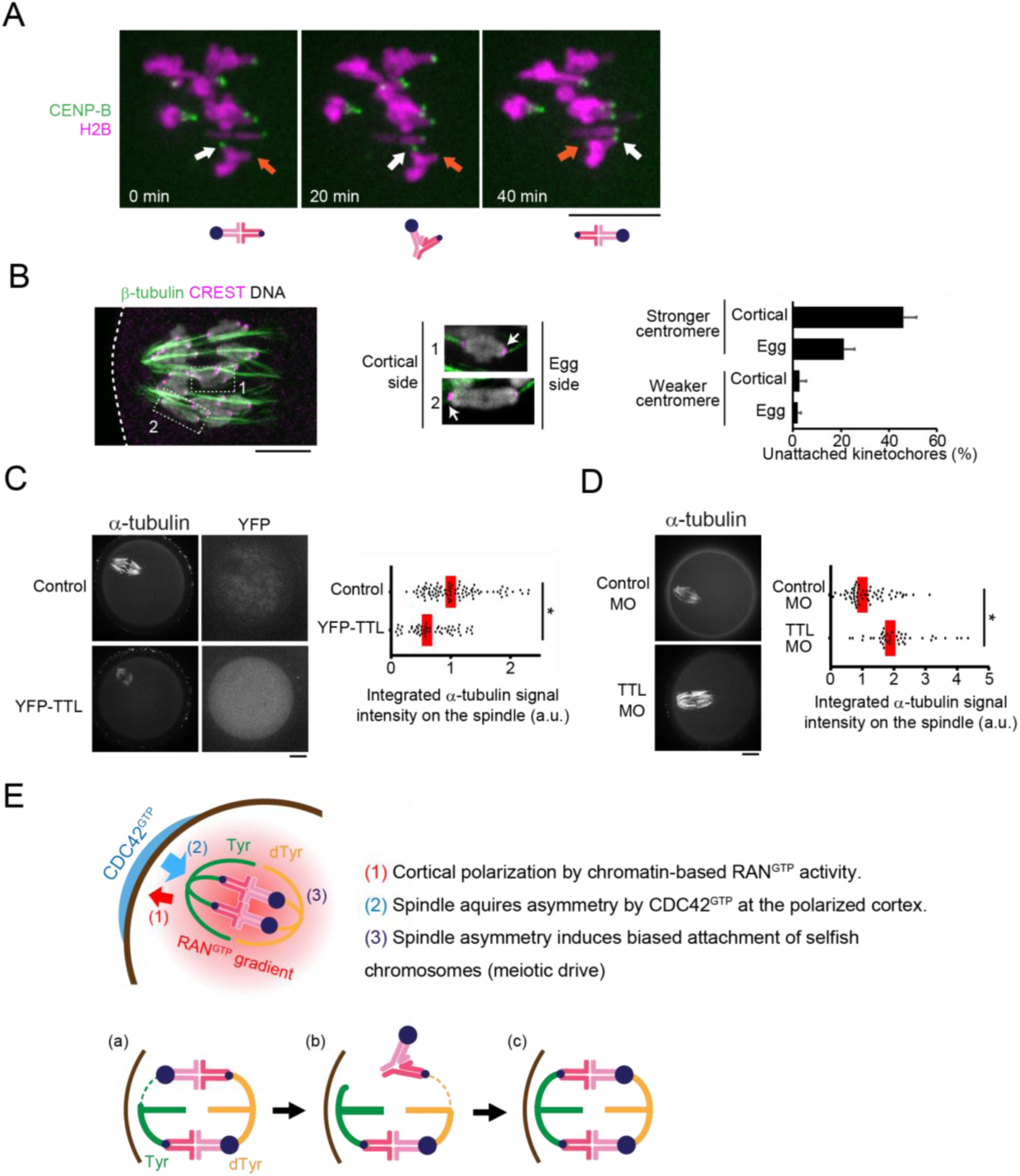
MT tyrosination promotes unstable interactions between selfish centromeres and the cortical side of the spindle. (**A**) CHPO x CF-1 oocytes expressing CENP-B-mCherry and H2B-EGFP were imaged live after spindle migration to the cortex (n = 23 cells). Time lapse images show an example of bivalent flipping; arrows indicate stronger (white) and weaker (orange) centromeres. (**B**) CHPO x CF-1 oocytes were fixed at 8 h after GVBD and analyzed for cold-stable MTs. Enlarged insets are optical slices showing individual bivalents with the stronger centromere (arrow) either facing the egg side and attached to cold-stable MTs (1) or facing the cortical side and not attached (2). Weaker centromeres are attached in both cases. Graph shows the percentage of centromeres without cold-stable attachments. Error bars represent s.d. for 3 independent experiments (> 50 bivalents analyzed in each experiment). (**C, D**) CF-1 oocytes expressing YFP-TTL or microinjected with morpholino against TTL were fixed at 6 h after GVBD and examined for cold-stable MTs. Graphs show integrated α-tubulin intensity in the spindle (n > 41 spindles for each sample). Each dot represents a single spindle; red line, median; *p < 0.0001. Images (A-D) are maximum intensity z-projections; scale bars, 10 μm. (**E**) Model for spindle asymmetry and meiotic drive. Top: cortical signals regulate MTs to induce tyrosination asymmetry within the spindle, and stronger centromeres (larger blue circles) attach preferentially to the egg side. Bottom: bivalent orientation is initially random (a), but attachments of stronger centromeres to the cortical side are unstable and tend to detach (b), leading to biased flipping to the egg side and biased orientation (c).

Our findings provide the first experimental evidence that asymmetry within the spindle is essential for meiotic drive and the first mechanistic insights into how signals from the cell cortex regulate MTs to induce spindle asymmetry and how selfish centromeres interact with the asymmetric spindle (Fig. 4E). Because the cortical side of the spindle will ultimately end up in the polar body, our findings explain how spindle asymmetry is consistently oriented relative to cell fate, providing spatial cues to guide the segregation of selfish elements. Moreover, the cortical signals are a product of cortical polarization, which is directed by chromosomes positioned near the cortex. This chromosome positioning is crucial for female meiosis because it allows the highly asymmetric division that is a universal feature of sexual reproduction in animals (*6*, *21*, *23*, *35*). Thus, selfish drive elements exploit the asymmetry inherent in female meiosis to bias their chances of transmission to the next generation.

## ACKNOWLEDGEMENTS

We thank BE. Black and MT. Levine for comments on the manuscript, G. Halet for the CDC42 and RAC constructs and discussions, R. Li for the RAN constructs, BL. Prosser for the detyrosinated α-tubulin and TTL antibodies. The research was supported by NIH grant GM107086 (M.A.L and R.M.S.) and by JSPS postdoctoral fellowship for research abroad and Research fellowship from Uehara Memorial Foundation (T.A.)

